# Early-life telomere length covaries with life-history traits and scales with chromosome length in birds

**DOI:** 10.1101/2021.08.07.455497

**Authors:** Michael Le Pepke, Thor Harald Ringsby, Dan T.A. Eisenberg

**Affiliations:** Centre for Biodiversity Dynamics (CBD), Department of Biology, Norwegian University of Science and Technology (NTNU), Trondheim, Norway; Department of Anthropology, University of Washington, Seattle, WA, USA; Center for Studies in Demography and Ecology, University of Washington, Seattle, WA, USA; Department of Biology, University of Washington, Seattle, WA, USA

**Keywords:** c-value, genome size, pace-of-life, phylogenetic comparative analysis, telomere biology, trade-offs

## Abstract

Telomeres, the short DNA sequences that protect chromosome ends, are an ancient molecular structure, which is highly conserved across most eukaryotes. Species differ in their telomere lengths, but the causes of this variation are not well understood. Here, we demonstrate that mean early-life telomere length is an evolutionary labile trait across 58 bird species (representing 35 families in 12 orders) with the greatest trait diversity found among passerines. Among these species, telomere length is significantly negatively associated with the fast-slow axis of life-history variation, suggesting that telomere length may have evolved to mediate trade-offs between physiological requirements underlying the diversity of pace-of-life strategies in birds. Curiously, within some species, larger individual chromosome size predicts longer telomere lengths on that chromosome, leading to the suggestion that telomere length also covaries with chromosome length across species. We show that longer mean chromosome length or genome size tends to be associated with longer mean early-life telomere length (measured across all chromosomes) within a phylogenetic framework constituting up to 32 bird species. Combined, our analyses generalize patterns previously found within a few species and provide potential adaptive explanations for the 10-fold variation in telomere lengths observed among birds.

## INTRODUCTION

Telomeres are an ancient molecular structure which is conserved across most eukaryotes (Fulnecková et al., 2013; Meyne, Ratliff, & Moyzis, 1989). Telomeres might have emerged when linear chromosomes evolved from a circular chromosome ancestor around 1.5 billion years ago (Lee, Leek, & Levine, 2017). Their function is to protect linear chromosomes during incomplete end-replication (Blackburn & Szostak, 1984) and oxidative stress (von Zglinicki, 2002), and in most tetrapods mean telomere length (TL) of somatic cells shortens through life (e.g. Tricola et al., 2018). Within some species, short telomeres have been shown to correlate with shorter lifespan (Heidinger et al., 2012), increased size (Ringsby et al., 2015), and oxidative stress (Reichert & Stier, 2017). Across species, a faster rate of telomere loss has been linked to shorter maximum species lifespan in several studies (Dantzer & Fletcher, 2015; Haussmann et al., 2003; Pepke & Eisenberg, 2020; Tricola et al., 2018; Vleck, Haussmann, & Vleck, 2003; Whittemore, Vera, Martínez-Nevado, Sanpera, & Blasco, 2019). However, no consistent patterns have emerged in how absolute TL is associated with lifespan or body mass across species (Gomes et al., 2011; Haussmann et al., 2003; Pepke & Eisenberg, 2021; Seluanov et al., 2007; Tricola et al., 2018).

The mean TL, measured across all chromosomes, has emerged as biologically relevant trait in evolutionary, ecological, and physiological studies (Monaghan, 2010; Nussey et al., 2014). Among wild birds, mean adult TL varies from around 5 kb in Western jackdaws (*Coloeus monedula*, Salomons et al., 2009) to more than 50 kb in great tits (*Parus major*, Tricola et al., 2018), but an evolutionary explanation for this 10-fold difference in mean TL is lacking (Tricola et al., 2018). A similar magnitude of variation is found within mammals (Gomes et al., 2011), in which TL is shorter in larger and longer-lived species (Pepke & Eisenberg, 2021). Furthermore, TL is positively associated with cancer risk across mammalian species (Pepke & Eisenberg, 2021). Thus, in mammals, evolution of shorter telomeres can be explained as an adaptation to counteract the increased risk of development of tumors associated with a larger number of cells and longer time to accumulate oncogenic mutations in larger and longer-lived species (Gomes et al., 2011; Gorbunova, Seluanov, Zhang, Gladyshev, & Vijg, 2014; Pepke & Eisenberg, 2021; Risques & Promislow, 2018; Seluanov et al., 2007; Tian et al., 2018). However, no association between TL and lifespan has been found in birds (Haussmann et al., 2003; Tricola et al., 2018; Vleck et al., 2003) and it is not known how TL covaries with body mass or other life-history traits in birds.

In life-history theory, evolutionary trade-offs are expected between vital life-history traits, such as between investment in current reproduction and investment in somatic growth, maintenance, or future reproduction (Stearns, 1989). Organisms can be placed along a fast-slow continuum of life-histories depending on how they resolve such trade-offs (Roff, 1993). A fast pace-of-life is characterized by higher investment in reproduction over survival, which is reflected in species with e.g. large clutch sizes and short generation times and lifespan (Araya-Ajoy et al., 2021; Ricklefs & Wikelski, 2002; Sæther, 1988). The variation in pace-of-life strategies is associated with physiological differences between species (Dammhahn, Dingemanse, Niemelä, & Réale, 2018; Ricklefs & Wikelski, 2002). We hypothesize that TL may have coevolved with suites of life-history traits and that TL may be used to rank species on the slow-fast axis of variation in life□history traits. Short telomeres could reflect decreased allocation of investment in somatic maintenance and hence increased allocation of resources to reproduction (Giraudeau, Angelier, & Sepp, 2019; Monaghan, 2010). However, long telomeres may be an adaptation to the cumulative negative effects of reproduction (Sudyka, 2019) and rapid growth on TL (Pepke et al., 2021), revealing the opposite relationship between TL and pace-of-life across species. We therefore investigate associations between TL and the slow-fast axis of life-history variation across species (Dantzer & Fletcher, 2015).

Curiously, within several species, cytogenetic studies have shown a positive correlation between the TL at a particular chromosome arm and the corresponding total chromosome length or chromosome arm length (reviewed in Klegarth & Eisenberg, 2018). This pattern has been found within laboratory mice (*Mus musculus*, Zijlmans et al., 1997), Chinese hamsters (*Cricetulus griseus*, Slijepcevic & Hande, 1999), humans (*Homo sapiens*, Graakjaer et al., 2003; Mayer et al., 2006; Suda et al., 2002; Wise et al., 2009), arabidopsis (*Arabidopsis thaliana*, Shakirov & Shippen, 2004), pearl millets (*Pennisetum glaucum*, Sridevi, Uma, Sivaramakrishnan, & Isola, 2002), yeast (*Saccharomyces cerevisiae*, Berthiau et al., 2006; Craven & Petes, 1999), and Tetrahymena (*Tetrahymena thermophila*, Jacob, Stout, & Price, 2004). Klegarth and Eisenberg (2018) tested whether this relationship extends across mammal species, using data on adult mean TL and mean chromosome length across 39 species of Primates and Cetartiodactyla (Gomes et al., 2011) and 11 species of Rodentia (Seluanov et al., 2007). They did not find any significant associations, but within Primates and Cetartiodactyla a positive trend between TL and chromosome length became stronger when the outlier Indian Muntjac (*Muntiacus muntjac*) was removed. Indian Muntjacs have recently undergone several whole chromosome fusions resulting in very large chromosomes (Wang & Lan, 2000). Perhaps an evolutionary lag accounts for their relatively short telomeres, which are adapted to shorter chromosomes (Klegarth & Eisenberg, 2018). Furthermore, many of the species included in past studies have been held in captivity or domesticated by humans, which might have altered their telomere length dynamics (Eisenberg, 2011; Manning, Crossland, Dewey, & Van Zant, 2002; Pepke & Eisenberg, 2021).

Birds may present an easier taxon with which to examine telomere–chromosome length co-evolutionary dynamics: In contrast to mammals, the avian karyotype is relatively conserved, and most birds have a chromosome number (2n) around 80 (Degrandi et al., 2020; Ellegren, 2010) suggesting that inter-chromosomal rearrangements are rare in birds. However, most bird species possess several microchromosomes, which have been shown to harbor functional, but ultra-long (“class III”) telomeres in some species (Atema, Mulder, van Noordwijk, & Verhulst, 2019; Delany, Gessaro, Rodrigue, & Daniels, 2007; Delany, Krupkin, & Miller, 2000; Nanda & Schmid, 1994; Nanda et al., 2002; Rodrigue, May, Famula, & Delany, 2005). Not all microchromosomes possess ultra-long telomeres (and not all species with microchromosomes possess any ultra-long telomeres, Delany et al., 2000; Nanda & Schmid, 1994; Nanda et al., 2002). Furthermore, these abnormal telomeres were found only on one chromosome arm and only in some individuals of inbred domestic chickens (*Gallus gallus*), whereas TL of the opposite chromosome arm was of typical size (“class II”, Delany et al., 2007). It is therefore possible that potential telomere–chromosome dynamics vary across chromosome size ranges (Atema et al., 2019), but this is unexplored in birds. Birds have the smallest genomes among extant amniotes, which is generally thought to represent adaptations to the metabolic requirements of active flight (Kapusta, Suh, & Feschotte, 2017; Wright, Gregory, & Witt, 2014). Yet, compared to mammals, some birds seem to have a larger amount of telomere sequences (Delany et al., 2000). Indeed, some of the ultra-long telomere signals may be attributed to sub-telomeric repeats (“class I”, Ingles & Deakin, 2016, but see Atema et al., 2019), which occur in some bird species (Nanda et al., 2002).

Here, we use mainly TL measurements of individuals of known age to obtain estimates of an early-life TL. We first reconstruct the evolution of mean TL of 58 bird species. We then investigate how TL relates to key life-history traits of birds. We then proceed to test the hypothesis that TL covaries with chromosome length across species. We do not resolve within-genome variation in TL across chromosomes, which is largely unknown in birds (Nanda et al., 2002), but we test whether variation in mean chromosome length underlies some of the variation observed in mean TL across species.

## MATERIALS AND METHODS

### Telomere length data

The use of methods to estimate relative amounts of telomeric DNA within samples (qPCR) limits the feasibility of comparative studies (Nussey et al., 2014). In this study we only used TL measured via the telomere restriction fragment (TRF) method (Haussmann & Vleck, 2002) or high-throughput quantitative fluorescence *in situ* hybridization (Q-FISH, Canela, Vera, Klatt, & Blasco, 2007; Lansdorp et al., 1996). In the TRF analysis, mean TL value is obtained from the distribution of TLs (in a Southern blot gel smear) across all chromosomes (Haussmann & Mauck, 2008). In the Q-FISH analysis, mean TL is obtained from the mean telomere probe fluorescence intensity across all chromosomes (quantified using microscopy image analysis, Canela et al., 2007). We searched the literature for avian telomere studies using TRF to measure mean TL from blood (Web of Science and Google Scholar [March 2021] search terms: “telomere”, “TRF”, “restriction fragment”, “Q-FISH”, “avian”, or “bird”). Correlations between TLs of various tissues suggest that blood TL is a good proxy of the TL across the organism (Daniali et al., 2013; Demanelis et al., 2020; Reichert, Criscuolo, Verinaud, Zahn, & Massemin, 2013). We used TL estimates from the same lab when available to minimize methodological effects (Tricola et al., 2018). Most studies used non-denatured in-gel TRF or Q-FISH to measure functional (terminal) TLs, except for 15 species indicated in Table S1 in the Supporting Information. These studies used denatured TRF, which may include short interstitial (class I) telomeric sequences, that may lead to underestimation of the mean functional TL (Foote, Vleck, & Vleck, 2013). The size distinction between class II and III telomeres is not well-defined (Atema et al., 2019; Delany et al., 2000) and some studies may not include the complete TL distribution if it is outside the range of the specific molecular size markers used (Atema et al., 2019; Foote et al., 2013; Haussmann & Mauck, 2008). However, since chromosome-specific TLs are largely unknown in birds, we rely on estimates of mean TL, which may reflect mainly “class II” telomeres.

To estimate an early-life TL for species with TL measurements at different reported ages, within each species we performed a linear regression of TL and individual age and used the extrapolated TL value at age 0 (i.e. the intercept, see Fig. 1 in Tricola et al., 2018). For these species, interspecific variance (97%) greatly exceeded intraspecific variance (3%) in TL, suggesting that our age correction method will only slightly change interspecific TL comparisons. For the remaining species, we used mean TL estimates for the youngest individuals reported (all <1 year old), except for 8 species (Table S1), where TL is averaged across unknown age classes. The domestic chicken was excluded because no mean TL has been reported (Delany, Daniels, Swanberg, & Taylor, 2003; Delany et al., 2000) and the chicken has a long history of human domestication, which is likely to have altered its telomere biology (Pepke & Eisenberg, 2021) and genome biology (Piégu et al., 2020). Variation in the activity of telomerase, a ribonucleoprotein capable of rebuilding telomeres (reviewed in Criscuolo, Smith, Zahn, Heidinger, & Haussmann, 2018), is unmeasured in our study and may confound the estimation of early-life TL.

**Fig. 1:**
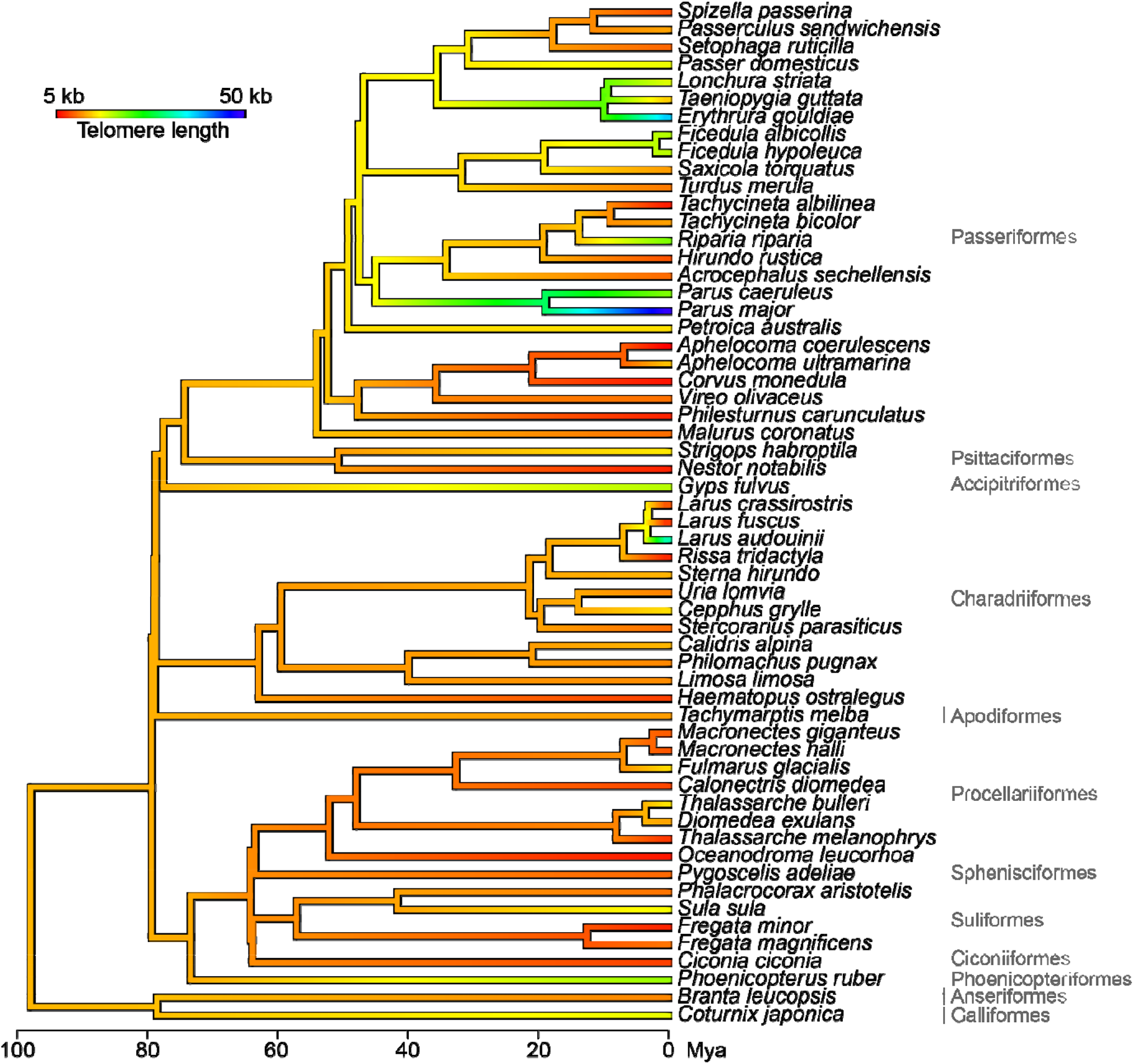
Calibrated maximum clade credibility tree illustrating the evolution of early-life telomere length in 58 bird species using a continuous color gradient (red: short telomeres; blue: long telomeres). Orders are shown on the right, and the timescale is in million years ago (Mya).

### Cytogenetic data

For 20 species estimates of genome size (the amount of DNA in a haploid genome, i.e. c-value) measured in erythrocytes were compiled from the Animal Genome Size Database, 2.0 (Gregory, 2020). When more than one c-value estimate was available in the database, we followed Gregory (2020) and averaged c-values across studies using up to four comparable cytometric methodologies (Hardie, Gregory, & Hebert, 2002, see Table S1). C-values (pg) were converted to mega base pairs (Mb) by multiplying with 978 Mb/pg (Doležel, Bartoš, Voglmayr, & Greilhuber, 2003). When these estimates were not available, we searched the National Center for Biotechnology Information genetic sequence database GenBank (Clark, Karsch-Mizrachi, Lipman, Ostell, & Sayers, 2016) and recorded the length of whole genome sequence (WGS) assemblies (in Mb, 10 species, in addition to 2 species from Grayson, Sin, Sackton, & Edwards, 2017, Table S1). Genome size estimates from cytometric methods are highly correlated with data obtained from WGS projects (Elliott & Gregory, 2015). Sequencing methods seem to underestimate genome size in some cases, a discrepancy that increases with absolute genome size (Elliott & Gregory, 2015). Since birds have relatively small genomes (1.0-2.2 Gb, Kapusta et al., 2017), we do not try to correct for this, but use cytometric estimates when available. Number of chromosomes was compiled from Degrandi et al. (2020). Atypical karyotypes are known from birds (Damas, O’Connor, Griffin, & Larkin, 2019; de Boer & van Brink, 1982) and we did not attempt to infer missing cytogenetic data from closely related species. Average chromosome lengths were calculated by first dividing genome size by the number of (haploid) chromosomes and then subtracting the telomeric DNA component from each chromosome (average TL multiplied by 2 representing the number of telomere arms per chromosome). For 14 species the karyotype is not yet known, and we therefore also test the association between TL and genome size, as a proxy for chromosome length across 32 species.

### Life-history data

Data on maximum lifespan (years) and average adult body mass (g) were compiled primarily from AnAge: The Animal Ageing and Longevity Database (Tacutu et al., 2018, Table S1 and S2), which combines data from captive and wild populations. Mean clutch size (mean number of eggs laid per clutch) and generation time (in years, based on modelled values of age at first reproduction, maximum lifespan, and annual adult survival) were primarily compiled from Bird et al. (2020), see Table S1. TL, mass, lifespan, and generation time were log_10_-transformed to linearize relationships observed in bivariate plots. We then used a phylogenetic principal component analysis (Revell, 2009) to construct a first principal component (PC1), which explained 55% of the variation in these traits and may reflect the fast (low values) to slow (high values) axis of variation in pace-of-life strategies (Table S3, Dantzer & Fletcher, 2015; Jeschke & Kokko, 2009).

### Phylogenetic reconstruction

We used the most recent time-calibrated avian phylogeny (Jetz, Thomas, Joy, Hartmann, & Mooers, 2012) based on the Hackett et al. (2008) backbone. We compiled a set of 1,000 trees from BirdTree.org and summarized these into a single maximum clade credibility tree using the maxCladeCred function in the ‘phangorn’ package in R (Schliep, 2011). This tree was pruned using the ‘ape’ package (Paradis & Schliep, 2018) and visualized using the ‘phytools’ package (Revell, 2012). Ancestral states were estimated using the function ‘fastAnc’ (Revell, 2012).

### Phylogenetic comparative analyses

Phylogenetic generalized least square regressions were performed using the ‘pgls’ function in ‘caper’ package (Orme et al., 2018), in which a variance-covariance matrix from the phylogenetic relationships (branch lengths) is compared to the actual covariance structure in the residual errors of the regression. The phylogenetic signal, *λ*, is a multiplier of the expected covariances (off-diagonal elements) that produces the actual variance-covariance matrix (Freckleton, Harvey, & Pagel, 2002; Pagel, 1999). Under the Pagel’s *λ* (PA) model, branch length transformations are optimized numerically using maximum likelihood within default bounds (0.0-1.0, Orme et al., 2018). When *λ*=0 the covariance between species is zero corresponding to a non-phylogenetic (ordinary) regression, or “star model” (ST), normally indicating that the traits are evolutionary very labile (Blomberg, Garland Jr, & Ives, 2003; Kamilar & Cooper, 2013). When *λ*=1 the evolution of the residual error is best approximated by a Brownian motion (BM) model of evolution (Felsenstein, 1985), which is the case for many gradually evolving traits (Kamilar & Cooper, 2013). The phylogenetic signal therefore estimates the extent to which associations between traits reflect their shared evolutionary history (Freckleton et al., 2002), i.e. the degree of similarity among closely related species compared to distantly related species. We ran bivariate linear regressions of log_10_-transformed TL (response variable) and chromosome length (18 species), genome size (32 species), PC1, log_10_-transformed maximum lifespan, and log_10_-transformed body mass (58 species), respectively. We also tested associations between TL and the life-history traits while accounting for either chromosome length or genome size using phylogenetic multiple regressions (Grafen, 1989). Due to the limited sample sizes, we ran alternative evolutionary models assuming either *λ*=1 (BM model), *λ*=0 (ST model) or the maximum□likelihood value of *λ* (PA model). We then used Akaike’s information criteria corrected for small sample sizes (AIC_c_) to determine the best model (Burnham & Anderson, 2002), which is reported in the results. All analyses were performed in R v. 3.6.3 (R Core Team, 2020).

### Sensitivity and outlier analyses

We performed sensitivity and outlier analyses within the phylogenetic context to test the robustness of our results to species sampling. We used a phylogenetic leave-one-out deletion analysis implemented in the ‘sensiPhy’ package (Paterno, Penone, & Werner, 2018) to test if any species are strongly affecting the associations. Species were sequentially removed one at a time and the phylogenetic regression refitted using ‘phylolm’ (Tung Ho & Ané, 2014). Highly influential species (outliers resulting in a standardized difference in *β*-estimates >2, Paterno et al., 2018) were then excluded and the regressions analyses rerun to obtain more robust phylogenetic regression estimates. We used a jackknifing method randomly removing a proportion of species (from 10 to 50%) and then refitted all regression models described above 500 times to estimate sensitivity of *β*-, *λ*-, and *p*-values to changes in sample size (Paterno et al., 2018).

## RESULTS

### Telomere length evolution

The greatest diversity in early-life TL was found within Passeriformes (6.2-50.5 kb, 25 species), whereas the other orders had relatively shorter and less variable TLs (7.1-34.9 kb, 33 species, Fig. 1). The ancestral TL of the 58 bird species was inferred, with a wide confidence interval, to be relatively short (12.4 kb, 95% confidence interval (CI)=[-1.1, 25.8], 97.9 Mya) compared to the range of TLs observed in extant species, but close to their average TL (14.0 kb). Phylogenetic signals in life-history traits were close to 1 (Table S4), but for TL *λ*=0.00 (CI=[0.00, 0.46]).

### Associations between telomere length and life-history traits

We found a significant negative association between TL and PC1 across all 58 species (best model: *β*_*PC1*_=-0.003±0.001 S.E., *p*=0.042, adjusted *R*^*2*^=0.055, *λ*=0.00, CI=[0.00, 0.27], Fig. 2a), suggestive of shorter TL in slower life-history species. There was also a negative association between TL and maximum lifespan (*β*_*log(lifespan)*_=-0.230±0.094 S.E., *p*=0.018, adjusted *R*^*2*^=0.080, *λ*=0.00, CI=[0.00, 0.27], Fig. 2b). Thus, a 1% increase in lifespan predicted a 0.23% decrease in TL. We found weak evidence for a negative association between body mass and TL (*β*_*log(mass)*_=-0.042±0.026 S.E., *p*=0.107, adjusted *R*^*2*^=0.029, *λ*=0.00, CI=[0.00, 0.28], Fig. 2c).

**Fig. 2:**
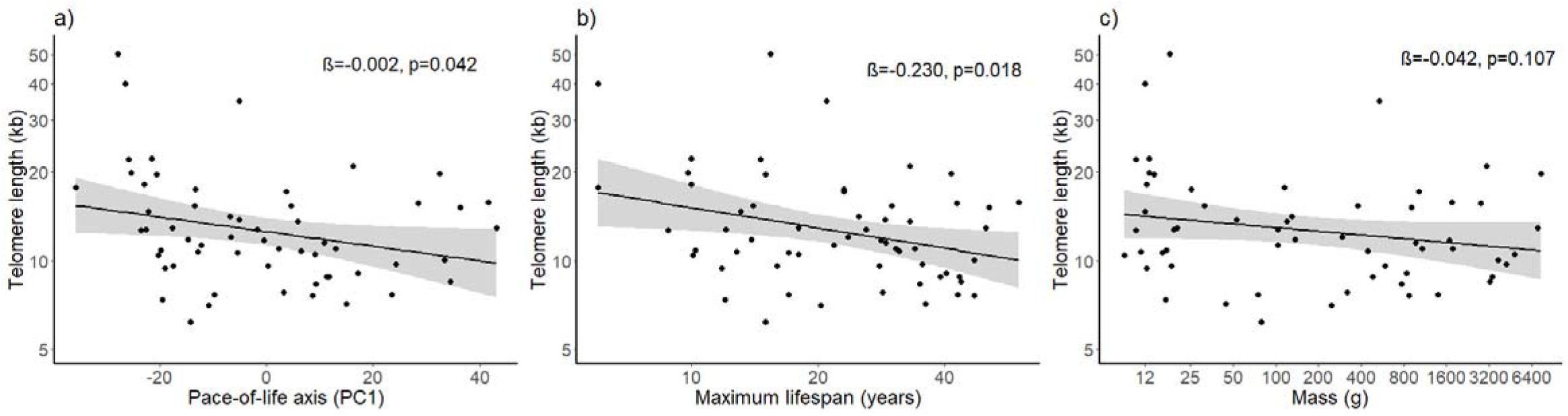
Associations between log_10_-transformed early-life telomere length (kb) and a) PC1 scores from a phylogenetic principal component analysis reflecting the slow-fast continuum of life-history trait variation across 58 bird species, b) log_10_-transformed maximum lifespan in years, and c) log_10_-transformed body mass in g. Scatter plots do not depict phylogenetic corrections. Phylogenetic regression lines and their associated statistics are shown. Grey shadings correspond to 95% confidence intervals.

We identified up to four phylogenetically significant outliers in the regressions between TL and life-history traits, but their exclusion only led to slight attenuations of the associations (see Figs. S1-S3 in the Supporting Information). These associations were generally robust to smaller sample size effects (mean change in *β* with 50% species removed was 36-52%) and the phylogenetic association was 0 in most simulations (Figs. S4-S6).

### Associations between telomere length, chromosome length and genome size

Genome size positively predicted chromosome length (*β*_*genome size*_=0.024±0.005 standard error [S.E.], *p*<0.001, adjusted *R*^*2*^=0.599, *λ*=1.00, CI=[0.00, 1.00], 18 species, Fig. S7), suggesting that genome size may be used as a proxy for chromosome length in birds.

We found weak evidence for a positive association between TL and chromosome length (best model: *β*_*log(chromosome length)*_=1.345±1.029 S.E., *p*=0.210, *R*^*2*^=0.210 *λ*=0.00, CI=[0.00, 1.00], 18 species, Fig. 3a). However, the phylogenetic outlier analysis identified one highly influential species, *Ciconia ciconia*, with a disproportional effect on the estimate (resulting in a change of *β* of 102%, see Paterno et al., 2018 and Fig. S8). Removal of this species revealed a substantial positive association between TL and chromosome length (*β*_*log(chromosome length)*_=2.710±1.106 S.E., *p*=0.027, *R*^*2*^=0.238 *λ*=0.00, CI=[0.00, 0.99], 17 species, Fig. 3a).

**Fig. 3:**
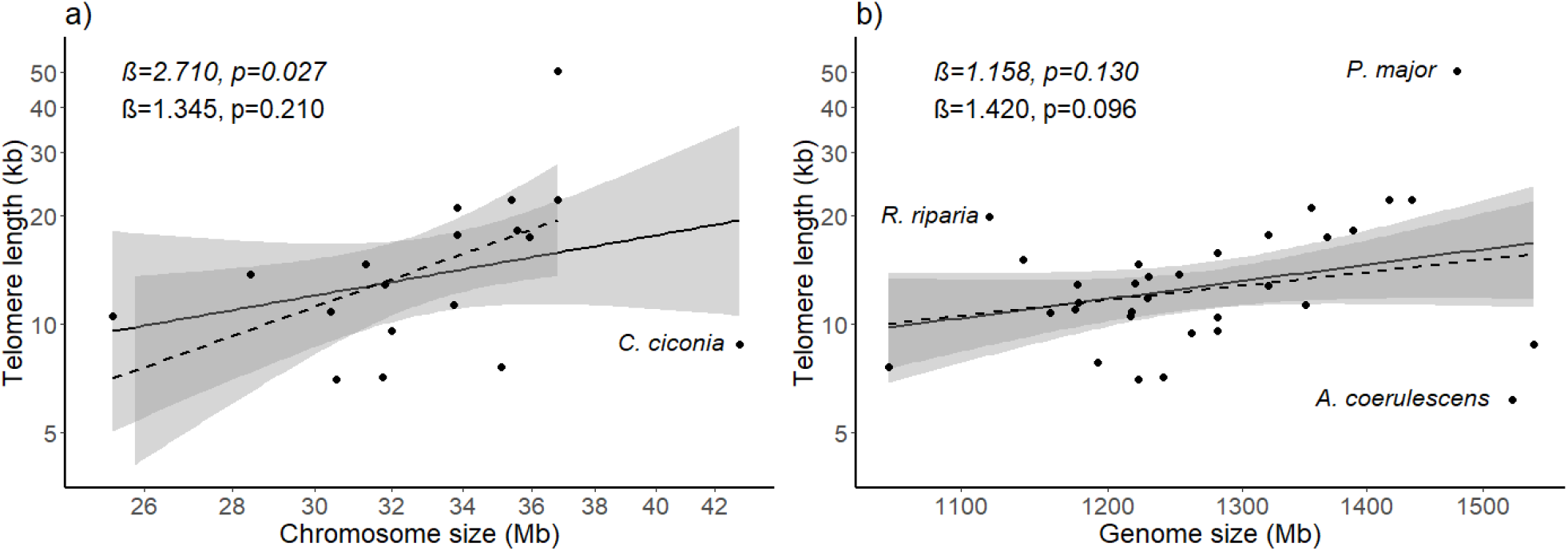
Associations between average log_10_-transformed early-life telomere length (kb) and a) chromosome size (Mb) for 18 bird species and b) genome size (Mb) for 32 bird species. Scatter plots do not show phylogenetic corrections. Phylogenetic regression lines and their associated statistics are shown (solid lines). The phylogenetically identified outliers *Ciconia Ciconia* (a), and *Aphelocoma coerulescens, Parus major*, and *Riparia riparia* (b) are excluded from the regression analyses shown in italics and with dashed regression lines. Grey shadings correspond to 95% confidence intervals.

There was a marginally significant positive association between TL and genome size (best model: *β*_*log(genome size)*_=1.420±0.826 S.E., *p*=0.096, adjusted *R*^*2*^=0.059, 32 species, Fig. 3b). We identified three highly influential species, *Aphelocoma coerulescens, Parus major*, and *Riparia riparia* (*β* changes of 50-61%, Fig. S9), the exclusion of which slightly attenuated the fit (*β*_*log(genome size*)_=1.158±0.742 S.E., *p*=0.130, *R*^*2*^=0.049, *λ*=0.00, CI=[0.00, 0.74], 29 species, Fig. 3b).

The associations between TL, chromosome length, and genome size were relatively unstable to larger reductions in sample sizes (changes in *β* with around 50% of the species removed were 45-81%, Fig. S10-S11).

### Multiple phylogenetic regressions of life-history and cytogenetic traits

The associations described above between life-history traits and TL, and between cytogenetic traits and TL were slightly or substantially attenuated when included in phylogenetic multiple regressions (Table S4). Mass, lifespan, and PC1 were only weakly correlated with genome size and chromosome length (Table S5), however, sample size was considerably reduced (by 44% when including genome size [32 species] and 68% when including chromosome length [18 species]).

## DISCUSSION

In this study, we showed that variation in mean early-life TL was significantly associated with key life-history traits underlying the pace-of-life continuum across 58 bird species. This result is consistent with the hypothesis that TL may be an important mediator of life-history trade-offs between reproduction, somatic maintenance, and cancer risk (Heidinger et al., 2012; Monaghan, 2010; Tian et al., 2018). Furthermore, we found some support for a positive association between TL and mean chromosome length or genome size. This indicates that a component of variation in TL may be constrained by interactions with chromosome length across species (Klegarth & Eisenberg, 2018; Slijepcevic, 2016).

Early-life TL was significantly negatively associated with maximum lifespan, but we only found weak evidence for a negative association with body mass (Fig. 2b-c). In mammals, lifespan and mass are strongly negatively associated with mean TL (Gomes et al., 2011; Pepke & Eisenberg, 2021), suggesting that these are general relationships across tetrapods. In birds, body size is constrained by adaptations to flight (Tobalske, 2016) and body masses within our study vary by almost 3 orders of magnitude compared to 7 orders of magnitude within the study of mammalian TL (Gomes et al., 2011), which may explain the weaker association between TL and mass in birds. However, there is still a large variation in TL particularly among small species, within both mammals and birds. This variation may be explained by the larger diversity of mechanisms evolved to deal with cancer defenses or cellular senescence within smaller bodied species (Risques & Promislow, 2018; Seluanov, Gladyshev, Vijg, & Gorbunova, 2018). That larger and longer-lived species of tetrapods have short telomeres, has been interpreted as an anti-cancer mechanism, limiting the risk of accumulating oncogenic mutations through replicative cell senescence (Campisi, 2001; Gorbunova et al., 2014; Pepke & Eisenberg, 2021). Research on cancer prevalence in wild bird species is very limited (Møller, Erritzøe, & Soler, 2017; Pesavento, Agnew, Keel, & Woolard, 2018). Boddy et al. (2020) found a positive relationship between litter size and cancer prevalence in 37 mammal species. They suggested that the faster pace-of-life associated with larger litter sizes, exposed fast life-history species to higher cancer rates by allocating more resources to offspring quantity than somatic maintenance. In our study, fast-lived species had a low PC1 score (i.e. short generation time and lifespan, large clutch size, and to some extent a small body mass, Table S3) and long telomeres compared to slow-lived species with a high PC1 score and short telomeres (Fig. 2a). If fast life-history bird species also have higher cancer rates, as suggested by Møller et al. (2017), these observations are consistent with the fact that longer telomeres are associated with increased cancer prevalence across species (in mammals, Pepke & Eisenberg, 2021). Thus, TL may have evolved to be longer to avoid the greater risk of critically short telomeres faced by species with accelerated TL shortening due to increased oxidative stress associated with high rates of reproduction (Sudyka, 2019). Selection for longer telomeres may have been further promoted by the lower antioxidant capacity or higher levels of oxidative damage found in bird species with a faster pace-of-life (Vágási et al., 2019; Xia & Møller, 2018). Bird species with a slower pace-of-life have also been found to have a lower telomere shortening rate (Dantzer & Fletcher, 2015), which suggests that TL and TL attrition co-vary across species, but this has not yet been shown (Tricola et al., 2018).

The sensitivity and outlier analyses indicated that the associations between TL and cytogenetic traits were susceptible to sample size effects (Figs. S8-S11). However, our results suggest an interaction between TL evolution and karyotype evolution. We found that a 1% increase in chromosome length was associated with a 2.7% increase in TL (Fig. 3a). The taxonomic diversity of species exhibiting positive scaling between TL and chromosome length within species (reviewed in the introduction) suggests that this is a highly conserved, fundamental characteristic of telomere biology. The explanation behind the positive correlation between telomere and chromosome lengths, remains unknown, but several molecular mechanisms may be involved (Klegarth & Eisenberg, 2018).

Experiments in yeast have shown that if telomeric and centromeric sequences are inserted into plasmids, they become unstable, probably because they are being pulled away from each other during mitosis (Enomoto, Longtine, & Berman, 1994). Slijepcevic (2016) suggested that this telomere–centromere antagonism could explain the correlation between TL and chromosome length observed within some species, i.e. the length of telomeres closer to centromeres is shorter to mitigate interference during mitosis. Furthermore, longer telomeres may be needed to protect longer chromosomes from end denaturation and rearrangements (Pampalona, Soler, Genescà, & Tusell, 2010; Slijepcevic, 1998). Supporting the connection between TL and chromosome size, Pontremoli et al. (2018) found that positive selection on genes implicated in telomere homeostasis among mammals was related to the number of chromosome arms. Given that genome size is relatively conserved among mammals (Kapusta et al., 2017), the positive selection at these genes is likely driven by chromosome size and these genes might help calibrate specific telomeres to the corresponding chromosomes. Assuming causality of the telomere– chromosome length association, more chromosome arms results in multiple short telomeres. This may facilitate chromosomal rearrangements (Murnane, 2012; Sánchez-Guillén et al., 2015; Slijepcevic, 1998), but also lead to a higher recombination rate (Pardo-Manuel de Villena & Sapienza, 2001).

Among mammals, the association between TL and chromosome length was highly influenced by the karyotypic abnormal Indian Muntjac (Klegarth & Eisenberg, 2018). However, the association remained non-significant after outlier exclusion. This study relied primarily on estimates of adult TL from cultured cell lines (Gomes et al., 2011). Our analysis may have better resolution by including mainly terminal TLs in early life, thereby reducing the effects of differing TL changes through life (Tricola et al., 2018). Consistent with the mammalian sensitivity to the karyotypic abnormal Indian Muntjac, our results were strongly influenced by the outlying white stork, *Ciconia ciconia* (8.8 kb, Fig. 3a), of a genus known to have undergone several chromosomal rearrangements (de Boer & van Brink, 1982, Fig. S7). For instance, *C. ciconia* (2n=72) probably has many more microchromosomes than the black stork, *C. nigra* (2n=52, de Boer & van Brink, 1982), whose mean TL we may then expect to be long, but that is currently unknown. The observation of ultra-long telomeres on some microchromosomes (Nanda et al., 2002) does not conform with the general patterns observed in this study. Since microchromosomes constitutes only around 23% of the avian genome size and are remarkably conserved across most bird species (Burt, 2002; O’Connor et al., 2019), it may be that the patterns reported here primarily retain to the telomere dynamics of macrochromosomes.

Within birds, larger genomes have been associated with lower metabolic rate (Vinogradov, 1997), reduced capacity for flight efficiency (Andrews, Mackenzie, & Gregory, 2009), and increased body size (Wright et al., 2014). The mechanism underlying these correlations may be acting through a positive relationship between genome size and cell size (Wright et al., 2014). However, if part of the variation in genome size is due to variation in telomere–chromosome length interactions, we suggest that some of these associations may involve adaptations in TL to different life-history strategies, as indicated in this study. For instance, correlations between life-history traits and genome size (Womack, Metz, & Hoke, 2019) may involve telomere–chromosome length dynamics.

We found TL to be evolutionary labile across bird species, as exemplified by the large intrageneric variation within *Aphelocoma, Larus, Tachycineta*, and *Thalassarche*, suggesting recent evolutionary change in TL (Fig. 1). Reconstructing the evolutionary history of TL changes within recent radiations of closely related species that represent independent replicated branching events, may elucidate adaptations underlying shifts in TL during speciation (Baird, 2018). As species progress through series of changes in species ecology and life-history (Pepke, Irestedt, Fjeldså, Rahbek, & Jønsson, 2019), associated changes in telomere biology may be observed within taxonomically more densely sampled clades (Canestrelli et al., 2020; Dupoué et al., 2017).

Our results indicate that some of the variation in early-life TL in birds arises through interactions with chromosome length, which may constrain the evolution of TL. Future cross-species studies attentive to the specificity of TL at different chromosome arms (Miga et al., 2020; Poon & Lansdorp, 2001), in particular of microchromosomes, may resolve the details of this interaction. Whether this effect has implications for telomere loss and the variation in senescence pattern across species remains unknown. However, mean TL also co-evolved with key life-history traits suggesting that the adaptive significance of TL may be as an important mediator of life-history trade-offs between investment in reproduction and somatic maintenance.

## ACKNOWLDGEMENT

We thank Mark Haussmann, Gianna Tricola, Simon Verhulst, Ellis Mulder, David S. Richardson, Antoine Stier, and Tiia Kärkkäinen for sharing data on telomere lengths within bird species, Claus Bech for providing longevity data, and Christina Bauch and Pat Monaghan for comments on earlier versions of this manuscript. This work is funded by the Research Council of Norway (Centre for Biodiversity Dynamics, 223257). The authors have no conflicts of interest to declare.

## AUTHOR CONTRIBUTIONS

MLP and DTAE conceived the ideas. MLP compiled and analyzed data and wrote the manuscript with contributions from all authors.

## DATA ACCESSIBILITY

All data is available from Table S1 in the Supporting information and from BirdTree.org, and it will be submitted to an open access data repository.

