## Supporting Information for "Early-life telomere length covaries with life-history traits and scales with chromosome length in birds"

### **Data and sources**

**Table S1:** Estimates of mean initial telomere length (TL), cytogenetic and life-history data gathered for this study. See Materials and Methods in the main text for main database sources (alternative sources are listed as footnotes). The taxonomy follows Gill et al. (2021). Dashes indicate data not available.

| Species name | Common name | Family | TL (kb) | Geno-me size (Mb) | Genome size method^(^^[1])^ | Chromo-some size (Mb) | Chromo-some number (n) | Body mass (g) | Max. lifespan (years) | Mean clutch  size | Genera-tion time (years) | PC1 | TL reference |
| --- | --- | --- | --- | --- | --- | --- | --- | --- | --- | --- | --- | --- | --- |
| *Acrocephalus sechellensis* | Seychelles warbler | Acrocepha-lidae | 10.6^(3)^ | - | - | - | - | 15.9 | 17.0 | 1.5 | 6.7 | -5.7 | Ellis Mulder & David S. Richardson, *unpublished*^(^^[2])^ |
| *Aphelocoma coerulescens* | Florida scrub jay | Corvidae | 6.2 | 1526 | FCM | - | - | 78.7 | 15.0 | 4.0 | 5.2 | -14.5 | Tricola et al., 2018 |
| *Aphelocoma ultramarina* | Transvolca-nic jay | Corvidae | 14.1 | - | - | - | - | 130.0 | 25.0 | 4.0 | 7.2 | -7.2 | Tricola et al., 2018 |
| *Branta leucopsis* | Barnacle goose | Anatidae | 11.7^(^^[3])^ | - | - | - | - | 1687.0 | 28.2 | 4.5 | 10.0 | -0.9 | Pauliny et al., 2012 |
| *Calidris alpina* | Dunlin | Scolopaci-dae | 13.7 | 1252 | FD | 28.5 | 44 | 52.5 | 28.8 | 4.0 | 7.2 | -5.4 | Pauliny et al., 2006 |
| *Calidris pugnax* | Ruff | Scolopaci-dae | 11.8 | 1229 | WGS | - | - | 136.0 | 13.9 | 4.0 | 5.2 | -14.9 | Tricola et al., 2018 |
| *Calonectris borealis* | Cory's shearwaters | Procellarii-dae | 9.1 | - | - | - | - | 836.9^(^^[4])^ | 40.3^(^^[5])^ | 1.0 | 13.7 | 16.6 | Bauch et al., 2020a |
| *Cepphus grylle* | Black guillemot | Alcidae | 15.4 | - | - | - | - | 378.0 | 29.9 | 1.5 | 9.2 | 4.2 | Tricola et al., 2018 |
| *Chloebia gouldiae* | Gouldian finch | Estrildidae | 40.0^(3)^ | - | - | - | - | 12.0 | 6.0 | 5.5 | 1.9 | -26.6 | Fragueira et al., 2019 |
| *Ciconia ciconia* | White stork | Ciconiidae | 8.8 | 1545 | FCM | 42.9 | 36 | 3350.0 | 39.0 | 4.0 | 12.5 | 11.2 | Pineda-Pampliega et al., 2020 |
| *Coloeus monedula* | Western jackdaw | Corvidae | 7.0 | 1223 | FD | 30.6 | 40 | 246.0 | 20.3 | 4.0 | 5.6 | -11.2 | Bauch et al., 2020b |
| *Coturnix japonica* | Japanese quail | Phasiani-dae | 17.7 | 1320 | FD, FCM | 33.9 | 39 | 115.0 | 6.0 | 9.5 | 2.2 | -35.9 | Stier et al., 2020 |
| *Cyanistes caeruleus* | Eurasian blue tit | Paridae | 22.1^(3)^ | 1438 | SCF | 36.9 | 39 | 10.3 | 14.6 | 7.5 | 2.9 | -26.1 | Atema et al., 2019 |
| *Diomedea exulans* | Wandering albatross | Diomedei-dae | 12.9^(13)^ | 1220 | WGS^(^^[6])^ | - | - | 7047.0 | 50.0 | 1.0 | 22.9 | 42.2 | Hall et al., 2004 |
| *Ficedula albicollis* | Collared flycatcher | Muscicapi-dae | 19.9 | 1118 | WGS | - | - | 12.7 | 9.8 | 6.0 | 2.3 | -25.5 | Stier et al., 2020 |
| *Ficedula hypoleuca* | European pied flycatcher | Muscicapi-dae | 19.7 | - | - | - | - | 13.9 | 15.0 | 6.0 | 4.1 | -20.8 | Kärkkäinen et al., 2019 |
| *Fregata magnificens* | Magnifi-cient frigatebird | Fregatidae | 11.0^(13)^ | 1177 | WGS | - | - | 1078.0 | 34.0 | 1.0 | 12.6 | 12.6 | Sebastiano et al., 2017 |
| *Fregata minor* | Great frigatebird | Fregatidae | 7.6 | - | - | - | - | 1400.0 | 43.0 | 1.0 | 17.2 | 22.9 | Tricola et al., 2018 |
| *Fulmarus glacialis* | Northern fulmar | Procellarii-dae | 15.1 | 1141 | WGS | - | - | 908.0 | 51.0 | 1.0 | 25.3 | 35.7 | Tricola et al., 2018 |
| *Gyps fulvus* | Griffon vulture | Accipitri-dae | 19.8^(^^[7])^ | - | - | - | 33 | 7436.0 | 41.4 | 1.0 | 17.0 | 31.6 | Whittemore et al., 2019 |
| *Haematopus ostralegus* | Eurasian oystercatcher | Haemato-podidae | 8.8 | - | - | - | 33 | 480.0 | 43.3 | 3.5 | 13.3 | 10.8 | Tricola et al., 2018 |
| *Hirundo rustica* | Barn swallow | Hirundini-dae | 9.6 | 1281 | FIA | 32.0 | 40 | 18.3 | 16.0 | 4.5 | 3.1 | -17.7 | Tricola et al., 2018 |
| *Larus audouinii* | Audouin's gull | Laridae | 34.9^(7)^ | - | - | - | - | 535.0 | 20.9 | 3.0 | 7.7 | -5.5 | Whittemore et al., 2019 |
| *Larus crassirostris* | Black-tailed gull | Laridae | 9.6 | - | - | - | - | 589.4^(^^[8])^ | 28.0^(8)^ | 2.5 | 8.2 | 0.0 | Mizutani et al., 2016 |
| *Larus fuscus* | Lesser black-backed gull | Laridae | 8.3^(13)^ | - | - | - | 34 | 766.2 | 34.9 | 2.5 | 12.6 | 8.9 | Foote et al., 2011a |
| *Limosa limosa* | Black-tailed godwit | Scolopaci-dae | 12.0^(3)^ | - | - | - | 45 | 294.0 | 23.6 | 4.0 | 7.7 | -7.0 | Atema et al., 2011; 2019 |
| *Lonchura striata* | White-rumped munia | Estrildidae | 18.2 | 1389 | FCM | 35.6 | 39 | 12.3 | 10.0 | 5.0 | 2.1 | -23.1 | Tricola et al., 2018 |
| *Macronectes giganteus* | Southern giant petrel | Procellarii-dae | 10.0^(13)^ | - | - | - | 40 | 3698.0 | 47.0 | 1.0 | 20.6 | 32.6 | Foote et al., 2011b |
| *Macronectes halli* | Northern giant petrel | Procellarii-dae | 9.7^(13)^ | - | - | - | - | 4206.3^(^^[9])^ | 35.4^(9)^ | 1.0 | 17.1 | 23.6 | Foote, 2008 |
| *Malurus coronatus* | Purple-crowned fairy-wren | Maluridae | 10.7^(3)^ | - | - | - | - | 11.1^(9)^ | 12.8^(9)^ | 2.5 | 4.2 | -13.1 | Eastwood et al., 2018 |
| *Nestor notabilis* | Kea | Strigopoi-dea | 7.6^(13)^ | 1054 | WGS | 35.1 | 30 | 867.5 | 47.0 | 3.0 | 7.9 | 8.1 | Horn, 2008 |
| *Oceanodroma leucorhoa* | Leach's storm petrel | Hydrobati-dae | 7.1 | 1240 | WGS^(6)^ | 31.8 | 39 | 44.6 | 36.0 | 1.0 | 14.8 | 14.5 | Tricola et al., 2018 |
| *Parus major* | Great tit | Paridae | 50.5 | 1477 | SCF | 36.9 | 40 | 17.9 | 15.4 | 8.5 | 3.0 | -28.0 | Tricola et al., 2018 |
| *Passer domesticus* | House sparrow | Passeridae | 17.5 | 1367 | FCM, FD, SCF | 36.0 | 38 | 25.3 | 23.0 | 4.5 | 3.7 | -13.6 | Ringsby et al., 2015 |
| *Passerculus sandwichensis* | Savannah sparrow | Emberizi-dae | 12.9 | 1178 | FIA | 31.9 | 37 | 20.2 | 18.0 | 4.5 | 2.2 | -17.8 | Tricola et al., 2018 |
| *Petroica australis* | South Island robin | Petroicidae | 15.4^(13)^ | - | - | - | - | 31.3^(^^[10])^ | 14.0^(^^[11])^ | 2.8^(^^[12])^ | 3.8 | -13.7 | Horn, 2008 |
| *Phalacrocorax aristotelis* | European shag | Phalacro-coracidae | 11.0^(13)^ | - | - | - | - | 1773.0 | 30.6 | 3.5 | 9.3 | 1.9 | Hall et al., 2004 |
| *Philesturnus carunculatus* | South Island saddleback | Callaeati-dae | 7.6^(13)^ | - | - | - | - | 74.7^(6)^ | 17.0^(9)^ | 2.0^(10)^ | 4.0 | -10.0 | Horn, 2008 |
| *Phoenicopte-rus ruber* | American flamingo | Phoenicop-teridae | 21.0^(7)^ | 1355 | FCM, FD | 33.9 | 40 | 3066 | 33.0^(9)^ | 1.0 | 13.0 | 15.7 | Whittemore et al., 2019 |
| *Pygoscelis adeliae* | Adélie penguin | Sphenisci-dae | 10.5 | 1217 | WGS | 25.3 | 48 | 4847.7^(10)^ | 18.0^(9)^ | 2.0 | 12.8 | 8.6 | Tricola et al., 2018 |
| *Rissa tridactyla* | Black-legged kittiwake | Laridae | 7.8^(3^^,^^[13])^ | 1193 | WGS | - | - | 317.0 | 28.5 | 2.0 | 9.8 | 2.9 | Schultner et al., 2014 |
| *Riparia riparia* | Sand martin | Hirundini-dae | 22.1 | 1418 | FIA | 35.5 | 40 | 12.7 | 10.0 | 4.5 | 2.3 | -21.7 | Pauliny et al., 2006 |
| *Saxicola torquatus* | African stonechat | Muscicapi-dae | 12.7 | - | - | - | 40 | 10.3 | 8.8 | 5.0 | 2.1 | -23.7 | Apfelbeck et al., 2019 |
| *Setophaga ruticilla* | American redstart | Parulidae | 10.4 | 1281 | FIA | - | - | 8.5 | 10.1 | 4.0 | 2.3 | -20.4 | Angelier et al., 2013 |
| *Spizella passerina* | Chipping sparrow | Passerelli-dae | 9.4 | 1262 | FIA | - | - | 12.2 | 11.8 | 4.0 | 2.5 | -19.3 | Foote et al., 2013 |
| *Stercorarius parasiticus* | Parasitic jaeger | Stercorarii-dae | 10.8^(13)^ | 1160 | WGS | - | - | 445.5 | 31.1 | 2.0 | 11.2 | 6.0 | Trondrud, 2017 |
| *Sterna hirundo* | Common tern | Laridae | 13.5 | 1230 | WGS | - | 34 | 120.0 | 33.0 | 2.0 | 10.4 | 5.5 | Tricola et al., 2018 |
| *Strigops habroptila* | Kakapo | Psittacidae | 15.8^(13)^ | 1281 | FIA | - | - | 1750.0 | 60.0 | 1.5 | 25.8 | 40.8 | Horn et al., 2011 |
| *Sula sula* | Red-footed booby | Sulidae | 17.1 | - | - | - | - | 1017.0 | 23.0 | 1.0 | 9.4 | 3.3 | Tricola et al., 2018 |
| *Tachycineta albilinea* | Mangrove swallow | Hirundini-dae | 7.4 | - | - | - | - | 16.7^(10)^ | 12.0^(^^[14])^ | 4.0 | 2.2 | -19.6 | Tricola et al., 2018 |
| *Tachycineta bicolor* | Tree swallow | Hirundini-dae | 12.8 | 1320 | FIA | - | - | 19.0 | 12.1 | 5.5 | 2.6 | -22.8 | Tricola et al., 2018 |
| *Tachymarpti melba* | Alpine swift | Apodifor-mes | 12.7^(3, 13)^ | - | - | - | - | 102.7 | 26.0 | 2.5 | 8.1 | -1.7 | Criscuolo et al., 2009 |
| *Taeniopygia guttata* | Zebra finch | Estrildidae | 14.7 | 1223 | FCM | 31.3 | 39 | 12.0 | 13.1^(^^[15])^ | 5.0 | 1.6 | -22.2 | Tricola et al., 2018 |
| *Thalassarche bulleri* | Buller’s albatross | Diomedei-dae | 15.6^(13)^ | - | - | - | - | 2781.5^(10)^ | 42.6^(^^[16])^ | 1.0 | 19.6 | 27.8 | Horn, 2008 |
| *Thalassarche melanophris* | Black-browed albatross | Diomedei-dae | 8.5^(13)^ | - | - | - | - | 3232.0 | 43.7 | 1.0 | 23.6 | 33.8 | Dupont et al., 2018 |
| *Turdus merula* | Common blackbird | Turdidae | 11.3 | 1350 | FD, SCF | 33.7 | 40 | 103.2 | 21.8 | 4.0 | 4.0 | -12.5 | Ibáñez-Álamo et al., 2018 |
| *Uria lomvia* | Thick-billed murre | Alcidae | 11.5 | 1179 | WGS | - | - | 964.0 | 29.0 | 1.0 | 12.9 | 10.3 | Tricola et al., 2018 |
| *Vireo olivaceus* | Red-eyed vireo | Vireonidae | 10.9 | 1218 | FIA | 30.4 | 40 | 17.0 | 10.2 | 4.0 | 2.6 | -20.0 | Foote et al., 2013 |

**New longevity record for Zebra finch**

**Table S2:** A colony of zebra finches (*Taeniopygia guttata*) was established by Claus Bech (*pers. comm.*) in January 2001 from individuals of reproductive age (3-4 months old) brought from a breeder. Housing conditions are described in Rønning et al. (2005) and all birds were provided with seed food and water *ad libitum* and allowed to breed.

| **ID** | **Sex** | **Hatched** | **Died** | **Lifespan (days)** | **Lifespan (years)** |
| --- | --- | --- | --- | --- | --- |
| Blue/blue/white/ring | Male | Around 2000/10/01 | 2013/11/01 | 4779 | 13.1 |
| 73 blue | Male | 2001/03/23 | 2014/02/27^(1)^ | 4724 | 12.9 |
| 156 striped | Male | 2001/06/14 | 2014/02/27^(1)^ | 4641 | 12.7 |
| ^(1)^ Last date the bird was observed alive. | | | | | |

**PCA of life-history trait variation**

**Table S3:** Loadings from a phylogenetically corrected principal component analysis (joint estimation of λ=0.86) of four life-history traits in 57 bird species. PC1 explained 55% of the variation among these traits. PC2 explained 22% of the variation and was primarily informed by body mass (89%).

| **Trait** | **PC1** | **PC2** |
| --- | --- | --- |
| Generation time | 0.914 | -0.126 |
| Maximum lifespan | 0.861 | -0.222 |
| Clutch size | -0.644 | 0.155 |
| Mass | 0.457 | 0.888 |

### **Phylogenetic outlier analyses of telomere length and life-history traits**

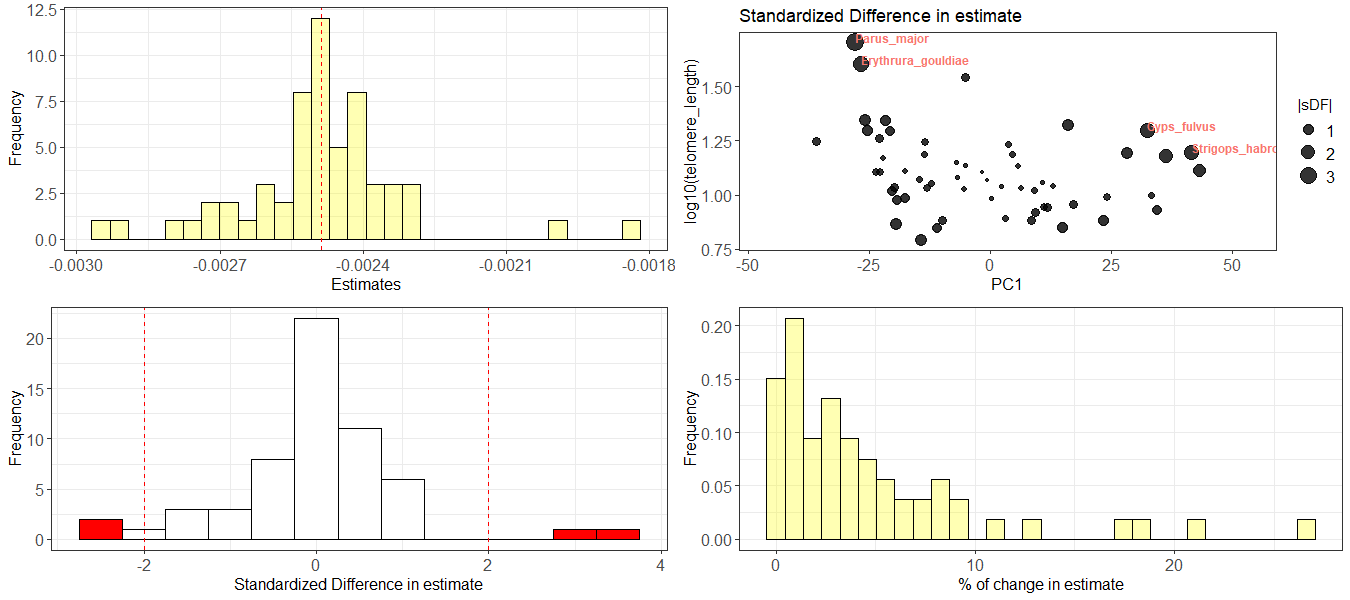
**Figure S1:** Outlier analysis of the phylogenetic regression between log_10_(TL) and PC1 (58 species, Fig. 2a). Four highly influential species (*Parus major*, *Chloebia gouldiae*, *Gyps fulvus*, *Strigops habroptila*) were identified. The association was slightly attenuated when these were removed (*β_PC1_* =-0.002±0.001 S.E., *p*=0.061, adjusted *R^2^*=0.048, *λ*=0.00, CI=[0.00, 0.23], 54 species).

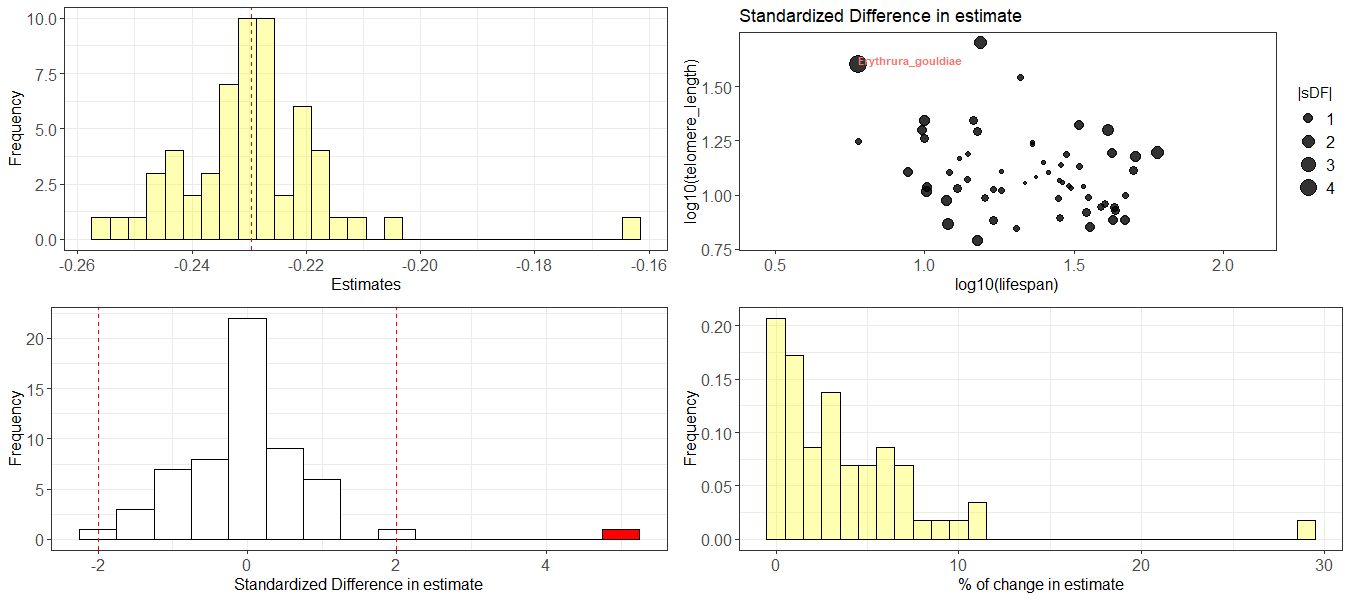
**Figure S2:** Outlier analysis of the phylogenetic regression between log_10_(TL) and log_10_(lifespan) (58 species, Fig. 2b). One highly influential species (*Chloebia gouldiae*) was identified. The association was attenuated when this species was removed (*β_log(lifespan)_*=-0.163±0.095 S.E., *p*=0.092, adjusted *R^2^*=0.033, *λ*=0.00, CI=[0.00, 0.25], 57 species).

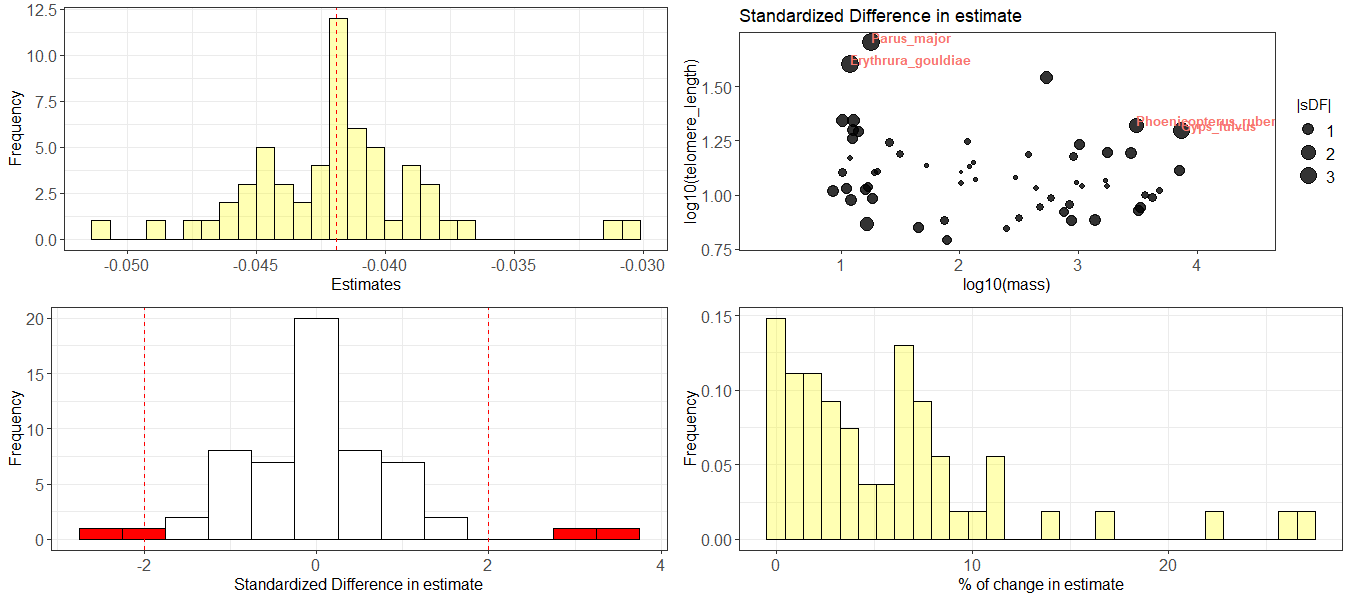
**Figure S3:** Outlier analysis of the phylogenetic regression between log_10_(TL) and log_10_(mass) (58 species, Fig. 2c). Four highly influential species (*Parus major*, *Chloebia gouldiae*, *Gyps fulvus*, *Phoenicopterus ruber*) were identified. The association was slightly attenuated when these were removed (*β_log(mass)_*=-0.035±0.022 S.E., *p*=0.119, adjusted *R^2^*=0.028, *λ*=0.00, CI=[0.00, 0.19], 54 species).

### **Phylogenetic sensitivity analyses of telomere length and life-history traits**

**Figure S4:** Sensitivity analysis of the phylogenetic regression between log_10_(TL) and PC1 (58 species, Fig. 2a). The *β*-estimates were relatively robust to sample size effects (mean change in *β* with 50% of the species removed was 42%).

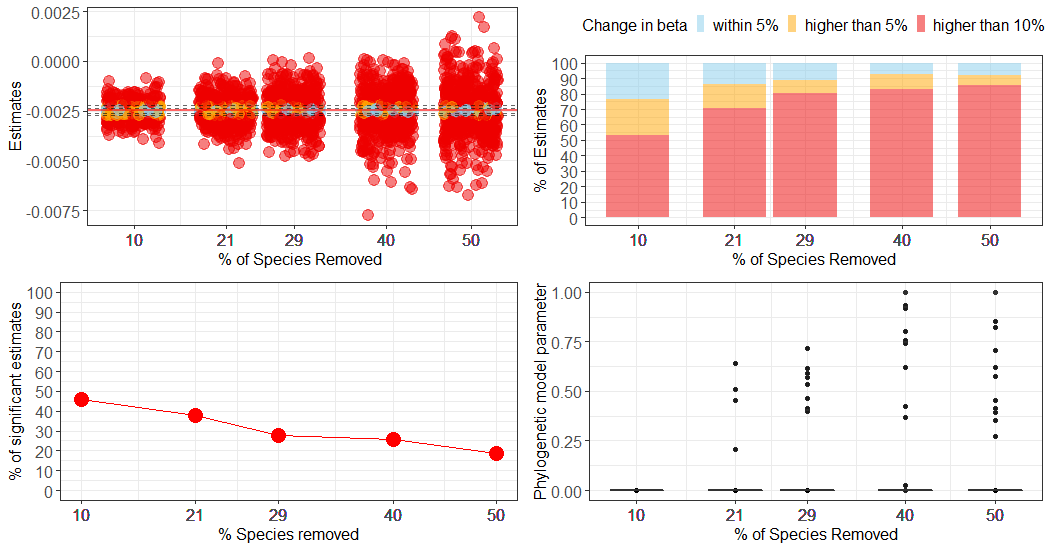

**Figure S5:** Sensitivity analysis of the phylogenetic regression between log_10_(TL) and log_10_(lifespan) (58 species, Fig. 2b). The *β*-estimates were relatively robust to sample size effects (mean change in *β* with 50% of the species removed was 36%), and 31% of the simulations suggested a significant relationship with 49% of the species removed.

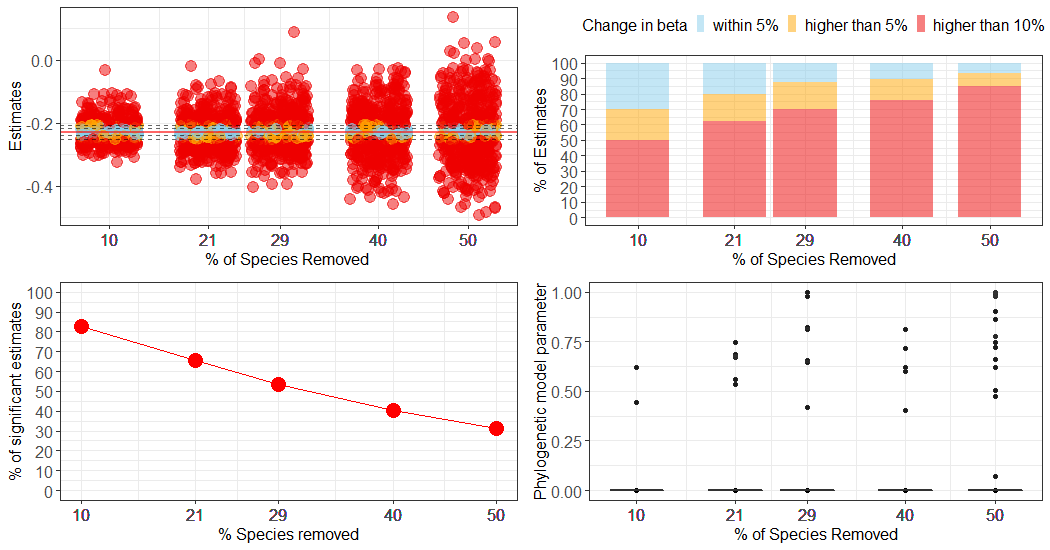

**Figure S6:** Sensitivity analysis of the phylogenetic regression between log_10_(TL) and log_10_(mass) (58 species, Fig. 2c). The *β*-estimates were relatively robust to sample size effects (mean change in *β* with 50% of the species removed was 52%). The association remained non-significant (p>0.05) under most simulations.

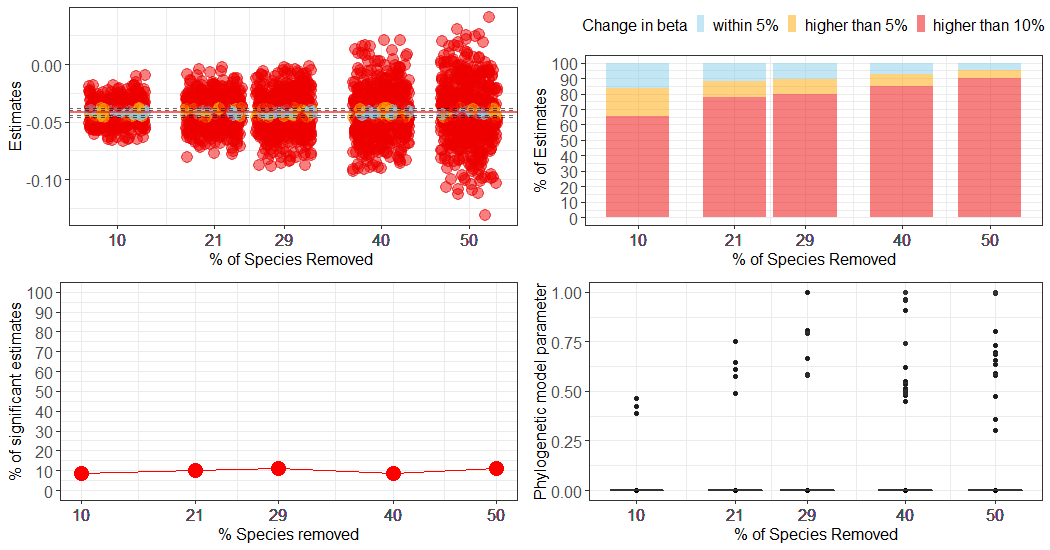

### **Association between chromosome length and genome size**

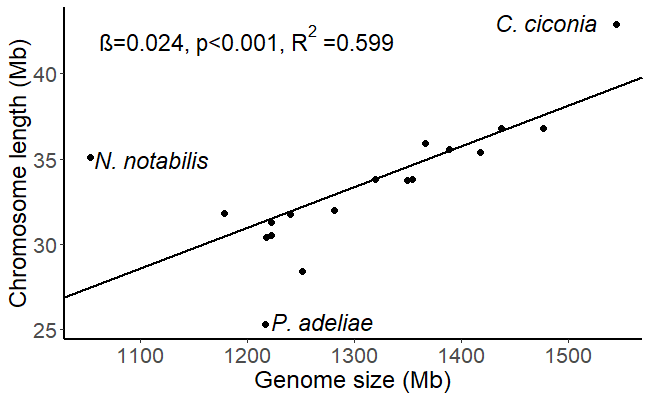

**Figure S7:** Genome size significantly predicted chromosome length in 18 bird species (*β_genome_* _size_=0.024±0.005 S.E., *p*<0.001, adjusted *R^2^*=0.599, *λ*=1.00, CI=[0.00, 1.00]). The outliers kea (*Nestor notabilis*, which has the smallest haploid chromosome number [30] and the smallest genome [1.05 Gb]), Adélie penguin (*Pygoscelis adeliae*), and white stork (*Ciconia ciconia*) are indicated, but not excluded from the analysis. Rearrangements of macrochromosomes have occurred within e.g. Psittaciformes and Ciconiiformes, resulting in an atypical chromosome numbers (de Boer & van Brink, 1982; Nanda et al., 2007).

### **Phylogenetic outlier analyses of telomere length and cytogenetic traits**

**Figure S8:** Outlier analysis of the phylogenetic regression between log_10_(TL) and log_10_(chromosome size) (18 species, Fig. 2a). One highly influential species (*Ciconia ciconia*) was identified.

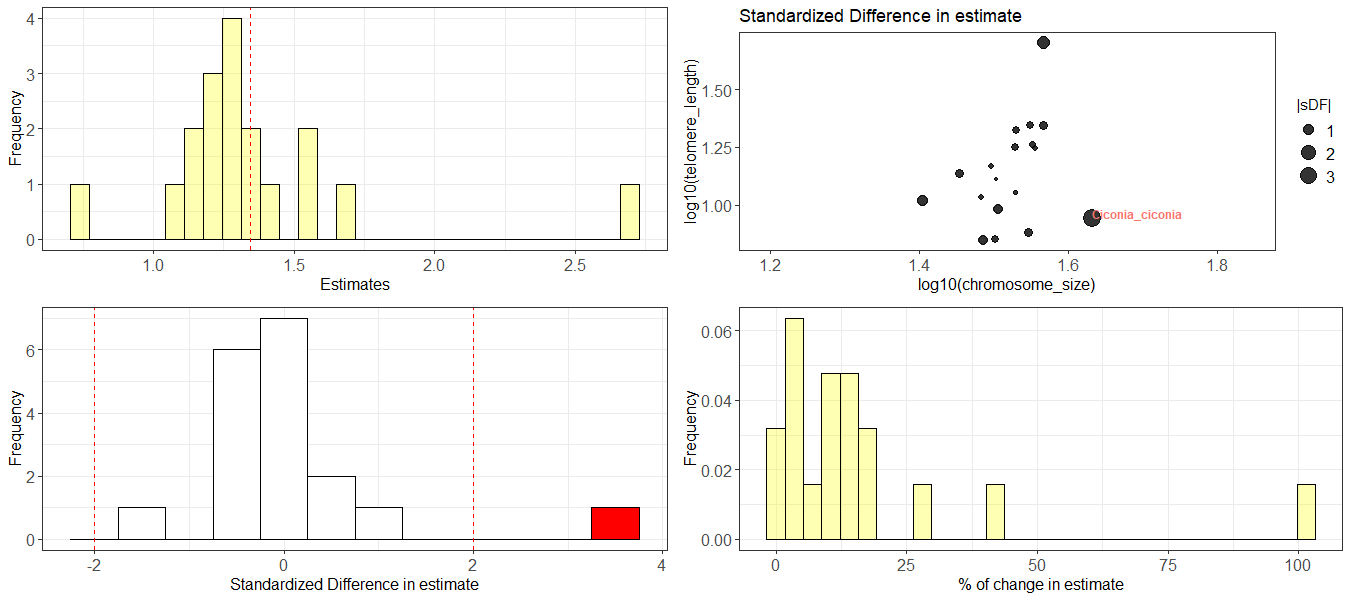

**Figure S9:** Outlier analysis of the phylogenetic regression between log_10_(TL) and log_10_(genome size) (32 species, Fig. 2b). Three highly influential species (*Parus major*, *Riparia riparia*, and *Aphelocoma coerulescens*^1^) were identified.

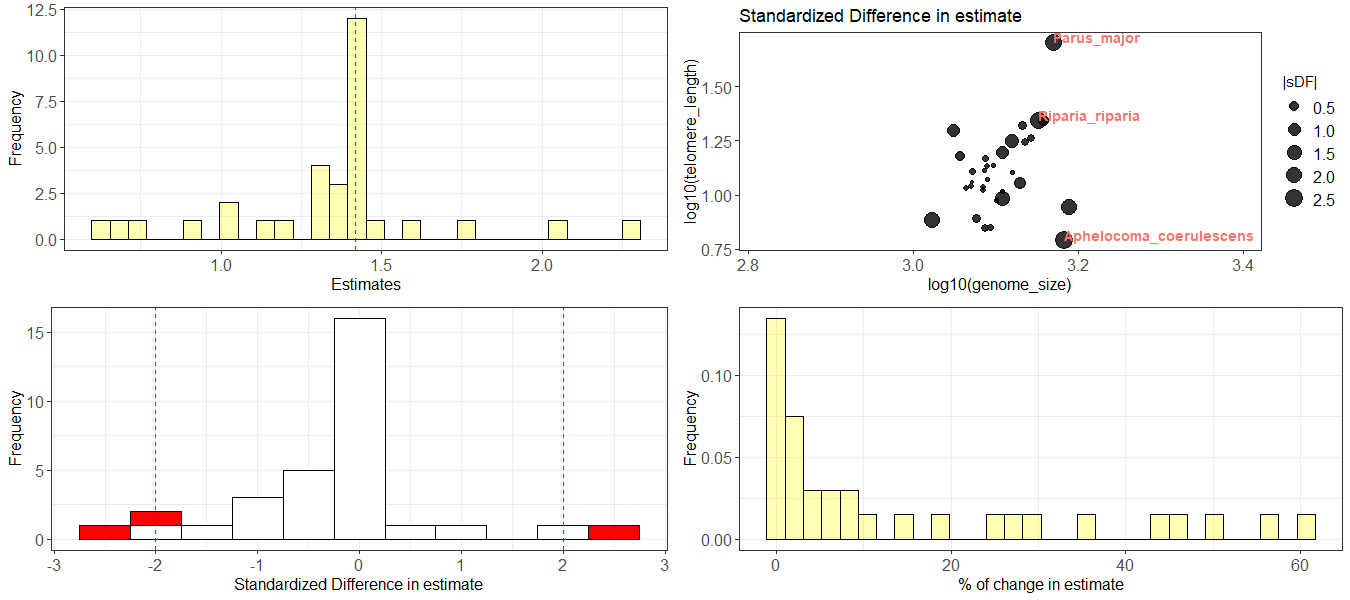

^1^ The Florida scrub jay (*A. coerulescens*) has very short mean TL (6.2 kb) despite a large genome (1.53 Gb; estimated using flow cytometry). However, all individuals (43) sampled within this species were older than 2 years and 40% were ≥7 years (see Tricola et al., 2018), which could lead to an underestimation of the extrapolated initial TL, which shortens fastest during early life in many bird species (e.g. Salomons et al., 2009). Furthermore, a genome assembly of *A. coerulescens* suggested that the flow cytometry genome size may be overestimated (WGS estimate of 1.08 Gb [Zhang et al., 2014], which is close to that of species with similarly short telomeres, e.g. *Nestor notabilis*, Table S1). In contrast, the congeneric larger transvolcanic jay (*A. ultramarine*) has telomeres twice as long (14.6 kb), but the genome size of this species has not yet been established.

### **Phylogenetic sensitivity analyses of telomere length and cytogenetic traits**

**Figure S10:** Sensitivity analysis of the phylogenetic regression between log_10_(TL) and log_10_(chromosome size) (17 species, Fig. 2a). The *β*-estimates were relatively robust with respect to small sample size effects (but mean change in *β* with 47% [8] of the species removed was 45% and the association remained significant [p<0.05] within 27% of the reduced datasets). Pagel’s *λ* (“phylogenetic model parameter”) collapsed at the boundary (0) for most simulations. The sample size is considerably reduced when reconstructing chromosome length evolution and power to detect phylogenetic signal in datasets with fewer than 20 species may be rather low (Freckleton et al., 2002; Blomberg et al., 2003; Garland et al., 2005) although Pagel’s *λ* have been shown to perform well in such case (i.e. resulting in a low rate of misidentification of phylogenetic signals in randomly evolving traits, Münkemüller et al., 2012).

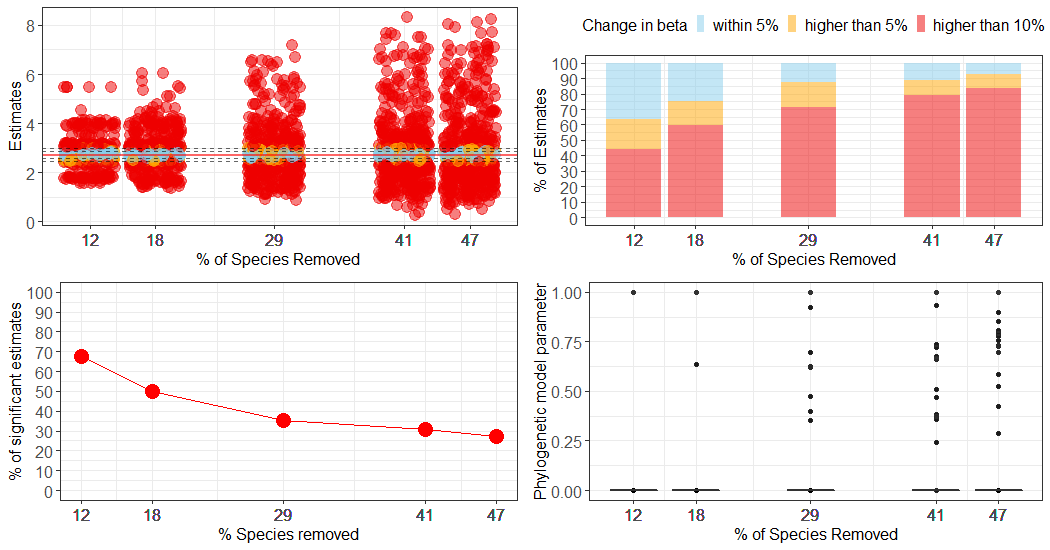

**Figure S11:** Sensitivity analysis of the phylogenetic regression between log_10_(TL) and log_10_(genome size) (29 species, Fig. 2b). The *β*-estimates were not robust to sample size effects (mean change in *β* with 48% [14] of the species removed was 81%).

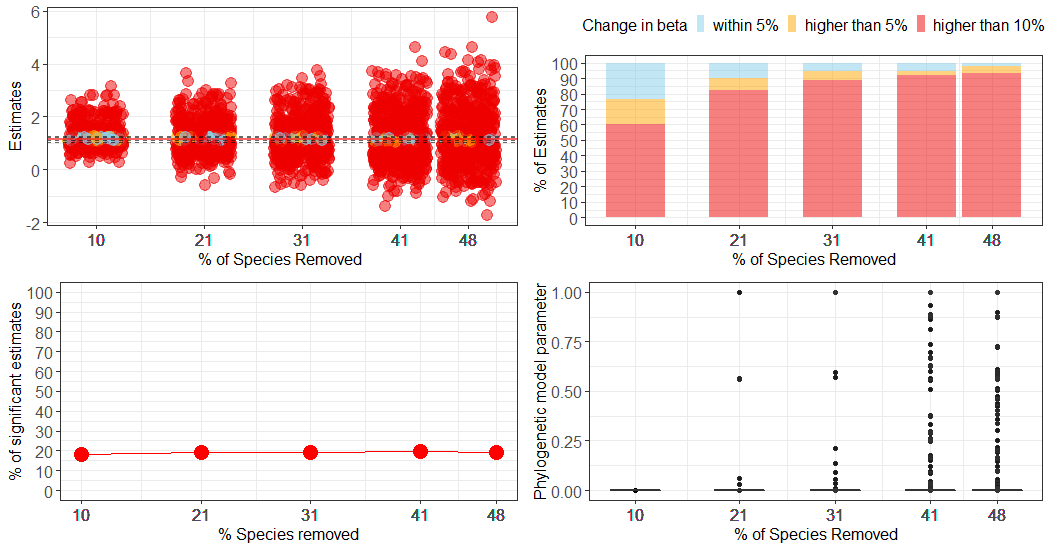

### **Multiple phylogenetic regressions of telomere length, cytogenetic and life-history traits**

**Table S4:** Phylogenetic multiple regression analyses of the associations between log_10_-transformed early-life telomere length (TL) as response variable, where A) body mass, B) maximum lifespan and C) PC1 are the focal explanatory variable within the total data set (58 species) and species subsets with data on genome size (32 species, second column) or chromosome length (18 species, third column). Phylogenetic signals in the log_10_-transformed species traits were respectively: TL (*λ*=0.00, CI=[0.00, 0.46]), chromosome length (*λ*=0.00, CI=[0.00, 1.00]), genome size (*λ*=0.22, CI=[0.00, 0.97]), body mass (*λ*=1.00, CI=[0.95, 1.00]), maximum lifespan (*λ*=0.86, CI=[0.65, 0.96]), and (untransformed) PC1 (*λ*=0.94, CI=[0.83, 0.99]). The bivariate associations between life-history traits and TL were strengthened in most cases when sample size was reduced from 58 to 18 species, but uncertainty of the regression coefficients increased, which is likely due to random sample size effects when the sample size becomes small (Figs. S4-6). The table also shows the estimates from phylogenetic multiple regressions controlling for genome size (32 species) or chromosome length (18 species). *β* denotes regression coefficients, *S.E.* denotes standard errors, *t* is the t-value, and *p* the significance value. *R^2^*-values reported are adjusted for the number of predictors.

| A | 58 species: log_10_(TL) | *β* | *S.E.* | *t* | *p* | 32 species: log_10_(TL) | *β* | *S.E.* | *t* | *p* | 18 species: log_10_(TL) | *β* | *S.E.* | *t* | *p* |
| --- | --- | --- | --- | --- | --- | --- | --- | --- | --- | --- | --- | --- | --- | --- | --- |
| intercept | | 1.099 | 0.024 | 46.641 | <0.001  *** | intercept | 1.096 | 0.033 | 33.069 | <0.001  *** | intercept | 1.060 | 0.055 | 19.347 | <0.001 *** |
| PC1 | | -0.003 | 0.001 | -2.084 | 0.042  * | PC1 | -0.002 | 0.002 | -1.372 | 0.180 | PC1 | -0.007 | 0.003 | -2.495 | 0.024* |
| *λ*=0.00 (95% CI: 0.00, 0.27), *R^2^*=0.055 | | | | | | *λ*=0.00 (95% CI: 0.00, 1.00), *R^2^*=0.028 | | | | | *λ*=0.00 (95% CI: 0.00, 0.91), *R^2^*=0.235 | | | | |
|  | | | | | | intercept | -2.540 | 2.709 | -0.938 | 0.356 | intercept | -0.607 | 1.385 | -0.438 | 0.668 |
|  |  |  |  |  |  | PC1 | -0.001 | 0.002 | -0.895 | 0.378 | PC1 | -0.007 | 0.003 | -2.377 | 0.031* |
|  |  |  |  |  |  | log_10_(genome size) | 1.172 | 0.874 | 1.342 | 0.190 | log_10_(chromo-some length) | 1.098 | 0.912 | 1.204 | 0.247 |
|  |  |  |  |  |  | *λ*=0.00 (95% CI: 0.00, 0.98), *R^2^*=0.053 | | | | | *λ*=0.00 (95% CI: 0.00, 0.97), *R^2^*=0.256 | | | | |
| B | **58 species: log_10_(TL)** | ***β*** | ***S.E.*** | ***t*** | ***p*** | **32 species: log_10_(TL)** | ***Β*** | ***S.E.*** | ***t*** | ***p*** | **18 species: log_10_(TL)** | ***β*** | ***S.E.*** | ***t*** | ***p*** |
| intercept | | 1.410 | 0.128 | 10.977 | <0.001  *** | intercept | 1.332 | 0.170 | 7.837 | <0.001  *** | intercept | 1.646 | 0.264 | 6.232 | <0.001  *** |
| log_10_(lifespan) | | -0.230 | 0.094 | -2.438 | 0.018* | log_10_(lifespan) | -0.172 | 0.127 | -1.351 | 0.187 | log_10_(lifespan) | -0.401 | 0.205 | -1.952 | 0.069^†^ |
| *λ*=0.00 (95% CI: 0.00, 0.27), *R^2^*=0.080 | | | | | | *λ*=0.00 (ST model), *R^2^*=0.026 | | | | | *λ*=0.00 (95% CI: 0.00, 1.00), *R^2^*=0.142 | | | | |
|  | | | | | | intercept | -2.450 | 2.727 | -0.898 | 0.376 | intercept | -0.673 | 1.409 | -0.478 | 0.640 |
|  |  |  |  |  |  | log_10_(lifespan) | -0.122 | 0.131 | -0.933 | 0.359 | log_10_(lifespan) | -0.434 | 0.196 | -2.217 | 0.042* |
|  |  |  |  |  |  | log_10_(genome size) | 1.197 | 0.861 | 1.389 | 0.175 | log_10_(chromo-some length) | 1.552 | 0.927 | 1.673 | 0.115 |
|  |  |  |  |  |  | *λ*=0.00 (95% CI: 0.00, 0.98), *R^2^*=0.055 | | | | | *λ*=0.00 (95% CI: 0.00, 1.00), *R^2^*=0.229 | | | | |
| C | **58 species: log_10_(TL)** | ***β*** | ***S.E.*** | ***t*** | ***p*** | **32 species: log_10_(TL)** | ***β*** | ***S.E.*** | ***t*** | ***p*** | **18 species: log_10_(TL)** | ***β*** | ***S.E.*** | ***t*** | ***p*** |
| intercept | | 1.196 | 0.062 | 19.284 | <0.001  *** | intercept | 1.210 | 0.080 | 15.186 | <0.001  *** | intercept | 1.305 | 0.115 | 11.346 | <0.001  *** |
| log_10_(mass) | | -0.042 | 0.026 | -1.639 | 0.107 | log_10_(mass) | -0.050 | 0.035 | -1.414 | 0.168 | log_10_(mass) | -0.088 | 0.055 | -1.600 | 0.129 |
| *λ*=0.00 (95% CI: 0.00, 0.28), *R^2^*=0.029 | | | | | | *λ*=0.00 (95% CI: 0.00, 1.00), *R^2^*=0.031 | | | | | *λ*=0.00 (95% CI: 0.00, 0.93), *R^2^*=0.084 | | | | |
|  | | | | | | intercept | -2.605 | 2.634 | -0.989 | 0.331 | intercept | -0.588 | 1.519 | -0.387 | 0.704 |
|  |  |  |  |  |  | log_10_(mass) | -0.039 | 0.036 | -1.094 | 0.283 | log_10_(mass) | -0.083 | 0.054 | -1.536 | 0.145 |
|  |  |  |  |  |  | log_10_(genome size) | 1.221 | 0.843 | 1.449 | 0.158 | log_10_(chromo-some length) | 1.238 | 0.991 | 1.250 | 0.231 |
|  |  |  |  |  |  | *λ*=0.00 (95% CI: 0.00, 0.97), *R^2^*=0.065 | | | | | *λ*=0.00 (95% CI: 0.00, 0.97), *R^2^*=0.115 | | | | |
| ^†^*p*<0.1, **p*<0.05, ***p*<0.01, ****p*<0.001 | | | | | | | | | | | | | | | |

**Correlation coefficients between explanatory variables**

**Table S5:** Pearson’s correlation coefficients between cytogenetic and life-history traits included in the multiple regressions in Table S2. We used the value of *λ*=0.00 (i.e. ST model) obtained in all association in Table S2.

|  | log_10_(mass) | log_10_(lifespan) | PC1 |
| --- | --- | --- | --- |
| 32 species: log_10_(genome size) | 0.120  (*p*=0.238) | 0.214  (*p*=0.214) | 0.265  (*p*=0.078) |
| 18 species: log_10_(chromosome length) | 0.239  (*p*=0.782) | 0.228  (*p*=0.692) | 0.221  (*p*=0.653) |
